# A single bout of mistimed eating is sufficient to cause lasting disruptions in the daily rhythms of metabolism and behavior

**DOI:** 10.64898/2026.09.22.753569

**Authors:** Siddhant Dharap, Maya Young, Erin N. Doherty, Lauren N Woodie

## Abstract

Synchrony between time-of-day and food intake is critical for maintaining homeostasis while desynchrony between these cues caused by repeated bouts of mistimed eating (ME) is profoundly disruptive to organismal health. However, it is unknown whether the same is true for a single bout of ME. While it is generally accepted that mammals recover quickly from acute alterations in feeding schedule and that chrono-disruption requires chronic, repeated changes, there have been no dedicated investigations of acute ME to suggest otherwise. Therefore, we exposed female and male C57Bl6/J mice to a single bout of ME or timed eating (TE) and continuously measured metabolic and behavioral responses for five days. A single bout of ME was sufficient to disrupt the daily rhythms of metabolic and behavioral outcomes for several days. Energy expenditure and wheel running activity were disrupted and light:dark variation in food intake and macronutrient utilization were altered for several days. Finally, ME changed the amount of time spent resting in for several days post-ME. The TE paradigm disrupted macronutrient utilization for two days but otherwise had no effect on metabolism or behavior. Our results demonstrate that one instance of ME alone has lasting impacts on daily rhythmicity and is an understudied source of circadian disruption.

Graphical Abstract

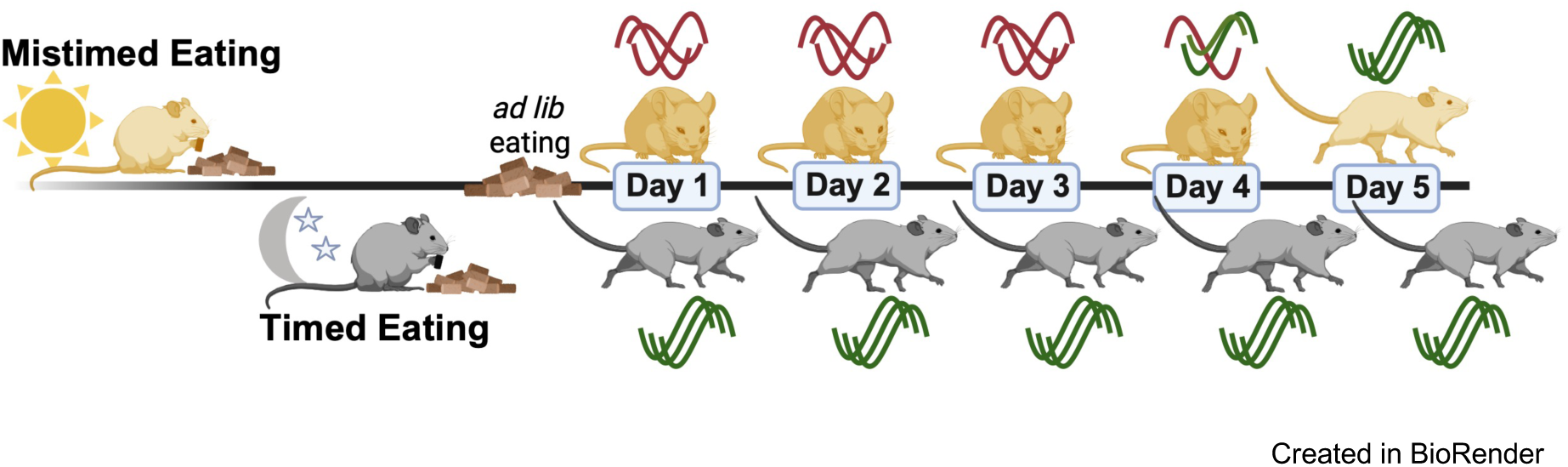

**NEW & NOTEWORTHY:** A single bout of mistimed eating (ME) produced lasting disruptions in the daily rhythms of metabolism and behavior. ME altered energy expenditure, activity, food intake, substrate utilization, and rest patterns for four days. ME effects manifested at different times in females and males highlighting a sex-specific vulnerability to the consequences of chrono-disruption. These findings suggest that a single bout of ME impairs metabolic flexibility and circadian behavior which are strongly implicated in human metabolic disease.

## INTRODUCTION

Mammalian physiology operates on an 24-hr clock, called circadian rhythms, that orchestrate the precise timing of biological processes ranging from metabolism to behavior (1,2). This oscillatory function requires coordinated integration of internal and external signals to maintain homeostasis of biological functions across the varying environmental states contained in a 24-hr day (2). At the molecular level, this is regulated by a transcription-translation feedback loop of clock genes that are rhythmically expressed (2,3). At the organismal level, many mammalian behaviors occur at certain times of day, such as when to sleep, work, or eat. Ultimately, the synchronization of external and internal oscillatory signals is what sets a healthy circadian rhythm (1–3).

The most commonly recognized circadian signal is light (4). Photic signals are processed by the master clock in the brain, the suprachiasmatic nucleus (SCN), which synchronizes the precise timing of hormonal, neural, and behavioral functions (4,5). However, light is not the only major timekeeper; food intake can also direct the timing of endogenous clocks (6). Tropic signals are detected by peripheral organs, such as the liver or gut, with downstream signaling modulating time-of-day specific tissue function and organ crosstalk (6–9). These major zeitgebers affect overlapping metabolic pathways, and thus, organismal health is heavily influenced by synchrony between the time of day and the timing of food intake (8,10).

As a result, individuals who often eat at the “wrong” time of day, prevalent in shift working professions (e.g., healthcare, transportation, manufacturing) are at greater risk for developing chronic disease conditions than those with a fixed, daytime working schedule (11). Recurrent mistimed eating (ME)-induced circadian disruptions are linked to exacerbated disease states such as cancer, mental health disorders, and metabolic diseases (12–15). However, it remains unclear whether an acute bout of ME is sufficient to induce biologically meaningful metabolic disturbances. This gap is significant as modern lifestyles require occasional late-night eating due to work demands, travel times, social obligations, and schedule variability (16–18). Thus, acute ME may be an unrecognized source of circadian disruption, with negative impacts on mammalian health and homeostasis. The goal of this study is to fill this knowledge gap and determine the extent to which a single bout of ME affects metabolic and behavioral outcomes.

## MATERIALS AND METHODS

### Animals

Six to eight-week-old, male and female C57Bl6/J mice were obtained from either Jackson Laboratories (Strain: #000664) or an in-house colony. Mice were provided with *ad libitum* access to PicoLab Rodent Diet 20 (5053, LabDiet) food and distilled water unless otherwise stated. Mice were housed in a 12:12 light:dark cycle at 22°C. All experimental procedures were approved by the George Washington University Institutional Animal Care and Use Committee and met the guidelines set out by the National Institutes of Health for the Care and Use of Laboratory Animals.

### Mistimed Eating and Timed Eating Paradigms

Experimental time was tracked by zeitgeber time (ZT) with ZT0 corresponding to light onset and ZT12 corresponding to light offset. To perform automated fasting and refeeding, animals were single-housed in Promethion metabolic cages (Sable Systems) outfitted with food hopper access control gates. Animals exposed to Mistimed or Timed Eating were first allowed to acclimate to the system for 24hrs (Figure 1; Day −4) before 48hrs of baseline metabolic and behavioral phenotyping data collection (Figure 1; Day −3 and Day −2).

**Figure 1.**
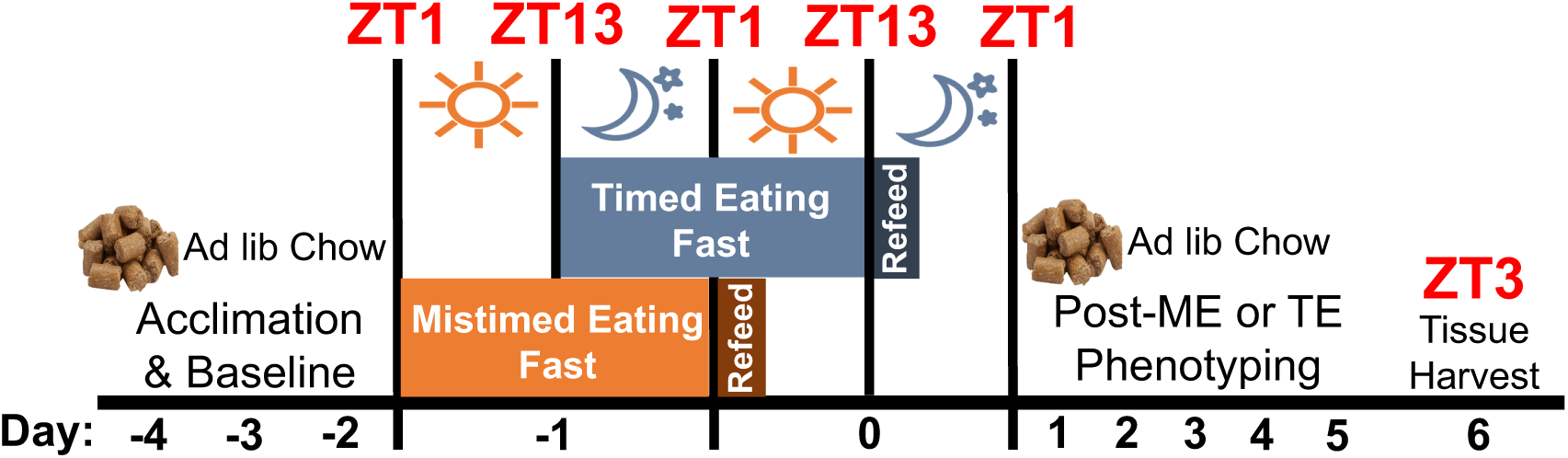
Timed and Mistimed Eating experimental paradigm and timeline. Experimental time was tracked by zeitgeber time (ZT) with ZT0 corresponding to lights on and ZT12 corresponding to lights off. Animals acclimated to the Promethion metabolic cages for one day (Day −4) before collection of baseline data for two days (Day −3 and −2) during which time they had *ad libitum* access to Chow and water. On Day −1 of the Mistimed Eating (ME) paradigm, animals were subjected to a 24hr fast beginning 1hr after lights on (ZT1). On Day 0, ME animals were automatically given access to their food hopper starting at ZT1 and animals were given 2hrs to consume food. On Day −1 of the Timed Eating (TE) paradigm, animals were subjected to a 24hr fast beginning 1hr after lights off (ZT13). On Day 0, TE animals were automatically given access to their food hopper starting at ZT13 and were given two hours to consume food. After Timed or Mistimed Eating animals remained in the metabolic cages for 5 days with *ad libitum* food access before tissue harvest at ZT3.

On Day −1 of the Mistimed Eating (ME) paradigm, animals were subjected to a 24hr fast beginning 1hr after light onset (ZT1) to avoid the potential confounding variable of light transitions affecting metabolism and behavior (Figure 1). On Day 0, ME animals were automatically given access to their food hopper starting at ZT1 (Figure 1). Food consumption was confirmed by observing real-time food intake data on Promethion Live software (Sable Systems), and the animals were given 2hrs to consume food from the time of their first eating bout. After 2hrs of mistimed eating, food access gates were closed until ZT6, after which time animals had *ad libitum* access to their food for the remainder of the experiment (Day 1 to Day 5). Animals that did not consume food during the refeed period were excluded from further analysis.

On Day −1 of the Timed Eating (TE) paradigm, animals were subjected to a 24hr fast beginning 1hr after light offset (ZT13) to avoid the potential confounding variable of light transitions affecting metabolism and behavior (Figure 1). On Day 0, TE animals were automatically given access to their food hopper starting at ZT13 (Figure 1). Food consumption was confirmed by observing real time food intake data on Promethion Live software (Sable Systems), and the animals were given 2hrs to consume food from the time of their first eating bout. After 2hrs of timed eating, food access gates were closed until ZT18, after which time animals had *ad libitum* access to their food for the remainder of the experiment (Day 1 to Day 5). Animals that did not consume food during the refeed period were excluded from further analysis.

### Behavioral and Metabolic Phenotyping

Behavioral and metabolic phenotype were determined by continuous measurement of energy expenditure (EE), macronutrient utilization (respiratory exchange ratio (RER)), food intake, non-wheel activity, wheel running, and time spent resting by the Promethion system. This data was analyzed in 1hr slices using Macro Interpreter software (Sable Systems). Data was further processed to create 6hr sums (EE, food intake, non-wheel activity, and wheel running) or averages (RER and time spent resting). Cycle data was determined by taking the sum (EE, food intake, non-wheel activity, and wheel running) or average (RER and time spent resting) over the entire light cycle or dark cycle. Total EE, food intake, non-wheel activity, and wheel running was determined by taking the sum over the light and dark cycle.

EE, via kCal, was determined using the Weir equation: EE=(3.941 x VO_2_ / 1.11 x VCO_2_) x 1.44. RER was determined by rate (mL/min) of carbon dioxide emission (VCO_2_) divided by the rate of oxygen consumption(VO_2_), with a ratio of 0.7 indicating lipid utilization and a ratio of 1.0 indicating carbohydrate utilization. Food intake was determined by detecting changes in food hopper mass with a standard deviation greater than 0.05g and minimum intake duration of 30 sec. General cage activity was determined by X and Y beam breaks excluding wheel running activity. Wheel running activity was separately determined by meters run on the wheel. Time spent resting was determined as a percentage of time during a 1hr slice that an animal spent not engaged in eating, drinking, grooming, or locomotion for 40sec or more.

### Postmortem Analyses

At the end of behavioral and metabolic phenotyping, the animals were euthanized at ZT3. We chose this time to assess the gross anatomical and physiologic effects of ME in recently fed animals while avoiding the potential confounding variable of light onset at ZT0. Trunk blood was collected and livers, inguinal white adipose tissue (iWAT), and gonadal white adipose tissue (gWAT) were excised and weighed.

Normalized weights for liver, iWAT, and gWAT were calculated by dividing tissue weight (g) over animal body weight (g). Non-fasting blood glucose was determined from trunk blood using a blood glucometer (ContourONE).

### Statistical Analyses

All data was graphed and analyzed using GraphPad Prism 11. Line graphs of metabolic and behavioral phenotyping data were analyzed using a two-way repeated measures ANOVA. When a significant interaction was detected, data was analyzed at the cycle level (light, dark, total) for each 24hr day. Bar graphs of metabolic phenotyping data were analyzed using a two-way ANOVA. When significance was detected in the ANOVA omnibus test, Tukey’s post-hoc test was used to determine differences among levels. ME and TE refeed food intake and postmortem measures were analyzed with a Mann-Whitney U test. Two-tailed statistical significance was determined at p<0.05. All data are presented as mean ± standard error measurement (SEM).

## RESULTS

### Timed and Mistimed Eating Groups were Similar at Baseline

To determine the metabolic and behavioral response to a single bout of mistimed eating, 6-8wk old C57Bl6/J female and male mice were randomly assigned to a Mistimed Eating (ME) protocol or a Timed Eating (TE) protocol (Figure 1). One female TE cage, one female ME cage, two male TE cages, and one male ME cage did not record wheel-running data and were excluded from wheel-running analysis but were included in all other analyses. Body weights were evenly distributed between the two groups (females *p*= 0.7996; Table 1, males: *p=* 0.2900; Table 2) and 48hrs of baseline measurements for EE, wheel running, food intake, RER, non-wheel activity, and time spent resting revealed no group differences prior to TE or ME (Supplementary Figure 1). One TE female, one ME female, two TE males, and four ME males did not consume food during the refeed period and were excluded from all further analyses. The remaining animals in TE and ME groups consumed the same amount of food during the refeed period (females: *p=* 0.7104; Figure 2A, males: *p=* 0.9999; Figure 2B).

**Figure 2.**
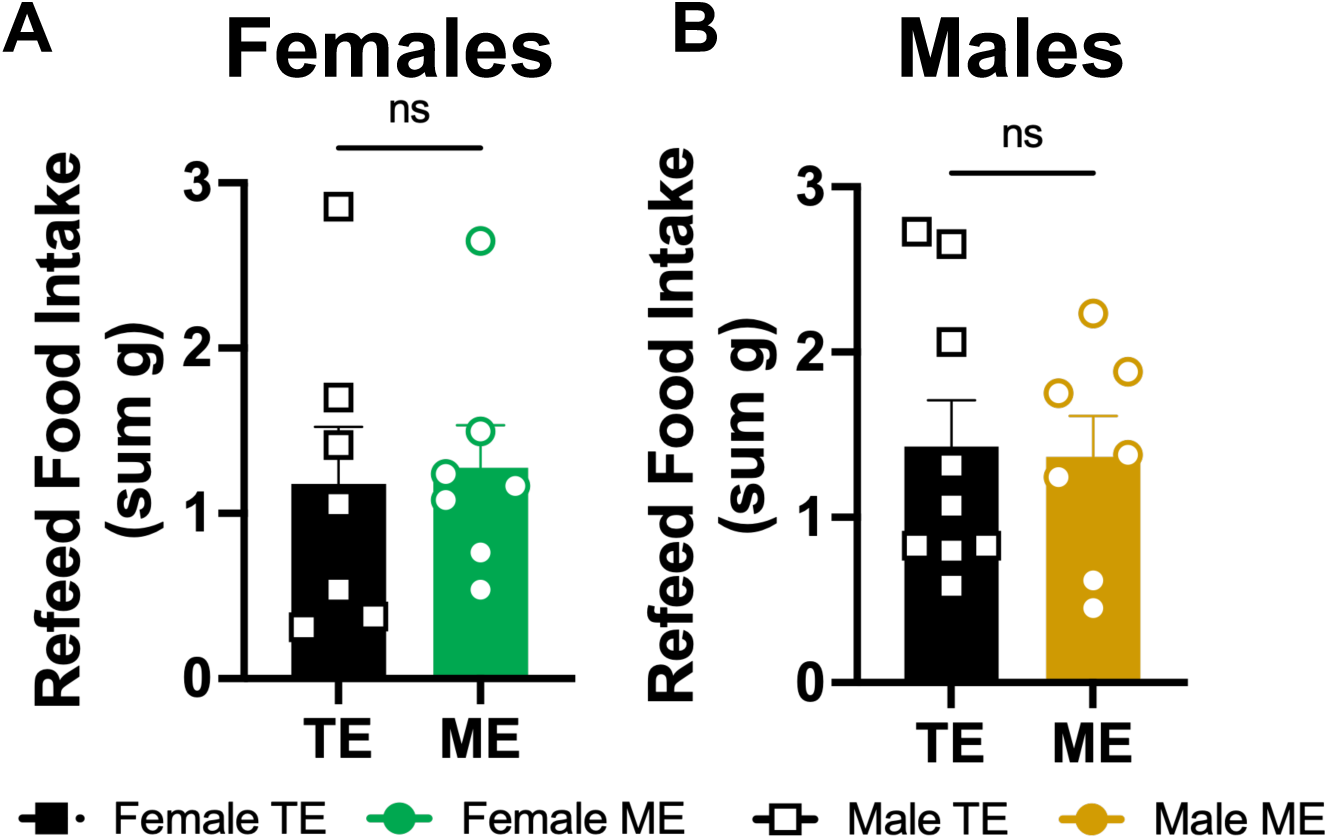
Animals consume the same amount of food in the timed and mistimed eating paradigms. (A) Sum of food consumed during the 2 hour refeed window in female animals exposed to the timed eating (TE) or mistimed eating (ME) paradigm. TE *n =* 7. ME *n =* 7. (B) Sum of food consumed during the 2 hour refeed window in male animals exposed to the TE or ME paradigm. TE *n =* 9. ME *n =* 7. Values are means ± SEM. Comparisons were made between ME vs. TE with a Mann-Whitney U test. ns= no significance.

**Table 1.**
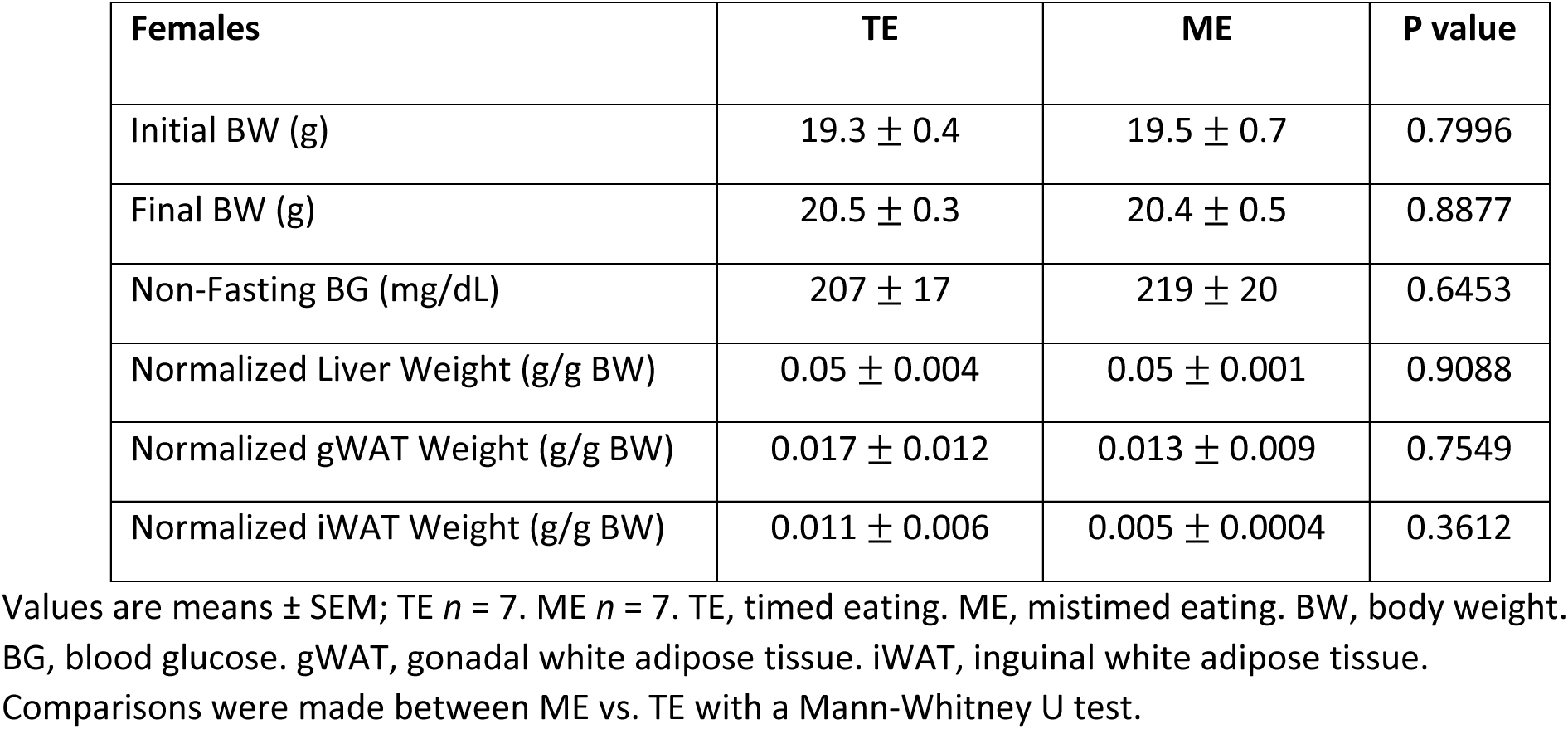
Female body weight and postmortem measurements.

| Females | TE | ME | P value |
| --- | --- | --- | --- |
| Initial BW (g) | 19.3 ± 0.4 | 19.5 ± 0.7 | 0.7996 |
| Final BW (g) | 20.5 ± 0.3 | 20.4 ± 0.5 | 0.8877 |
| Non-Fasting BG (mg/dL) | 207 ± 17 | 219 ± 20 | 0.6453 |
| Normalized Liver Weight (g/g BW) | 0.05 ± 0.004 | 0.05 ± 0.001 | 0.9088 |
| Normalized gWAT Weight (g/g BW) | 0.017 ± 0.012 | 0.013 ± 0.009 | 0.7549 |
| Normalized iWAT Weight (g/g BW) | 0.011 ± 0.006 | 0.005 ± 0.0004 | 0.3612 |

Values are means ± SEM; TE *n* = 7. ME *n* = 7. TE, timed eating. ME, mistimed eating. BW, body weight. BG, blood glucose. gWAT, gonadal white adipose tissue. iWAT, inguinal white adipose tissue. Comparisons were made between ME vs. TE with a Mann-Whitney U test.

**Table 2.** Male body weight and postmortem measurements.

| Males | TE | ME | P value |
| --- | --- | --- | --- |
| Initial BW (g) | 24.5 ± 0.9 | 23.9 ± 0.6 | 0.5498 |
| Final BW (g) | 25.4 ± 1.0 | 24.3 ± 0.8 | 0.3823 |
| Non-Fasting BG (mg/dL) | 232 ± 17 | 192 ± 16 | 0.0920 |
| Normalized Liver Weight (g/g BW) | 0.06 ± 0.001 | 0.05 ± 0.002 | 0.6367 |
| Normalized gWAT Weight (g/g BW) | 0.008 ± 0.0005 | 0.007 ± 0.0008 | 0.0938 |
| Normalized iWAT Weight (g/g BW) | 0.007 ± 0.0005 | 0.006 ± 0.0009 | 0.6696 |

Values are means ± SEM; TE *n* = 7. ME *n* = 4. TE, timed eating. ME, mistimed eating. BW, body weight. BG, blood glucose. gWAT, gonadal white adipose tissue. iWAT, inguinal white adipose tissue. Comparisons were made between ME vs. TE with a Mann-Whitney U test.

### A Single Bout of Mistimed Eating Disrupts Patterns of Energy Expenditure

ME reduced energy expenditure during the 5-day measurement period (female Time x Group: *p*< 0.0001, Figure 3A; males Time x Group: *p=* 0.0251, Figure 3B). Upon further inspection at the cycle level, we found that the effects of ME manifested two days after the eating bout in female animals. On Day 3 and 4, ME reduced dark phase EE (female Day 3: *p=* 0.0043, Figure 3C *left panel*; female Day 4: *p=* 0.0039, Figure 3C *right panel*) and total EE (female Day 3: *p=* 0.0037, Figure 3C *left panel*; female Day 4: *p=* 0.0101, Figure 3C *right panel*) although light:dark variation remained intact.

**Figure 3.**
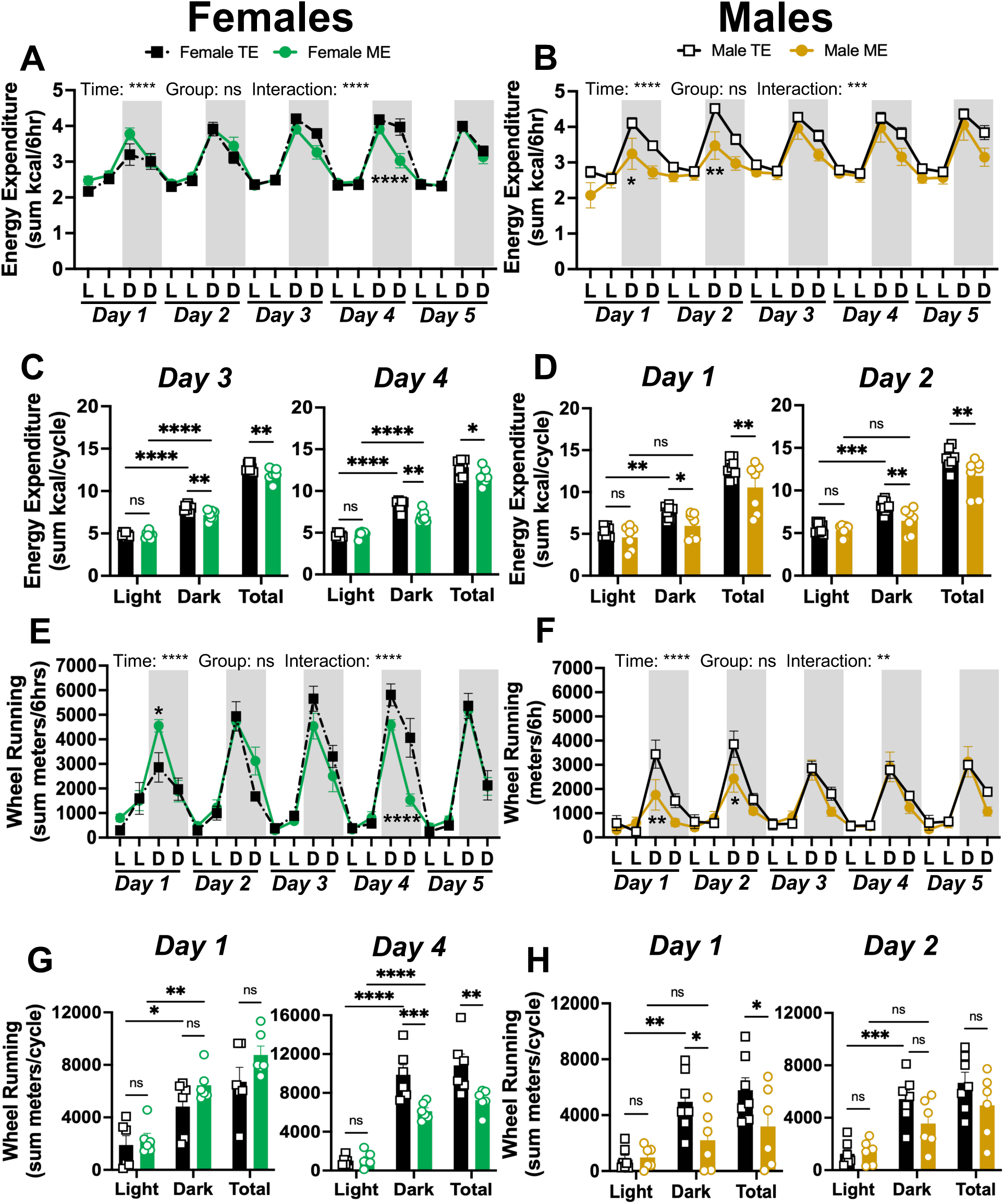
A single bout of mistimed eating disrupts the daily patterns of energy expenditure and wheel-running. (A) Sum of energy expenditure every 6 hours across the light (L) and dark (D) cycles for 5 days after timed eating (TE) or mistimed eating (ME) in female animals. TE *n* = 7. ME *n =* 7. (B) Sum of energy expenditure every 6 hours across the L and D cycles for 5 days after TE or ME in male animals. TE *n* = 9. ME *n =* 7. (C) The sum of energy expended during the light cycle, dark cycle, and total over 24 hours in female mice 3 (*left*) and 4 (*right*) days after TE or ME. TE *n* = 7. ME *n =* 7. (D) The sum of energy expended during the light cycle, dark cycle, and total over 24 hours in female mice 1 (*left*) and 2 (*right*) days after TE or ME. TE *n =* 9. ME *n* = 7. (E) Sum of wheel-running every 6 hours across the L and D cycles for 5 days after TE or ME in female animals. TE *n* = 7. ME *n =* 7. (F) Average of RER every 6 hours across the L and D cycles for 5 days after TE or ME in male animals. TE *n* = 7. ME *n =* 6. (G) The sum meters run on the wheel during the light and dark cycle over 24 hours in female mice 1 (*left*) and 4 (*right*) days after TE or ME. TE *n* = 7. ME *n =* 7. (H) The sum of meters run on the wheel during the light and dark cycle over 24 hours in female mice 1 (*left*) and 2 (*right*) days after TE or ME. TE *n =* 7. ME *n* = 6. Values are means ± SEM. In Figures 3A, 3B, 3E, and 3F significance was determined by repeated measures 2-way ANOVA with Holm-Šídák’s post hoc test. In Figures 3C, 3D, 3G, and 3F, significance was determined by 2-way ANOVA with Tukey’s post hoc test. ns= no significance, *p<0.05, **p<0.01, ***p<0.001, ****p<0.0001.

Males exhibited an earlier and exaggerated effect of ME. On Day 1 and 2, ME reduced dark phase EE (males Day 1: *p=* 0.0394, Figure 3D *left panel*; males Day 2: *p=* 0.0055, Figure 3D *right panel*) and abolished light:dark variation in EE (males Day 1: *p=* 0.2093, Figure 3D *left panel;* males Day 2: *p=* 0.1562, Figure 3D *right panel*). ME did not reduce light:dark variation in EE in female animals (Figure 3C; Supplementary Figure 2A), but male animals did not recover daily variation in EE until Day 3 post-ME (Supplementary Figure 2B). Female and male TE animals retained daily patterns in EE for the entire 5-day observation period (Figure 3C-D; Supplementary Figure 2A-B).

The ME-induced decrease in EE was partially driven by a reduction in wheel running activity (female Time x Group: p< 0.0001, Figure 3E; males Time x Group: *p=* 0.0460, Figure 3F). A reduction in wheel-running did not manifest in female animals until Day 4 when ME reduced dark phase wheel-running (female Day 4: *p=* 0.0007, Figure 3G *right panel*) and total wheel-running in females (female Day 4: *p=* 0.0011, Figure 3G *right panel*). Interestingly, ME caused females to wheel-run more in the early dark phase of Day 1 (female Day 1 1^st^ D: *p=* 0.0312; Figure 3E), but this did not result in an increase in dark cycle wheel-running on Day 1 (Figure 3G).

ME impacted male wheel-running patterns for the two days immediately following. On Day 1, ME reduced dark phase wheel running (male Day 1: *p=* 0.0181, Figure 3H *left panel*) and total wheel running activity (male Day 1: *p=* 0.0244, Figure 3H *left panel*). ME reduced light:dark variation in wheel running activity on both Day 1 and Day 2 (Figure 3H *left* and *right panels*). ME did not reduce light:dark variation in wheel-running in female animals (Figure 3G; Supplementary Figure 2C). However, male ME animals did not recover daily variation in wheel running until Day 3 post-ME (Supplementary Figure 2D *left panel*). Female and male TE animals retained daily patterns in wheel running activity for the entire 5-day observation period (Figure 3E-H; Supplementary Figure 2D-E).

### A Single Bout of Mistimed Eating Alters Food Intake and Substrate Utilization Patterns

Female animals exposed to a single bout of ME exhibited disrupted food intake patterns in the days immediately following ME, contrary to the delayed phenotype observed in EE and wheel-running (females Time x Group: *p=* 0.0107, Figure 4A). ME abolished light:dark variation in food intake during Day 1 (females Day 1: *p=* 0.2988, Figure 4C *left panel*), Day 2 (females Day 2: *p=* 0.5675, Figure 4C *right panel*), and Day 3 (females Day 3: *p=* 0.0530, Supplementary Figure 3A *left panel*), but did not impact the amount of food consumed (Figure 4C; Supplementary Figure 3A). Interestingly, both ME and TE females failed to exhibit daily variation in food intake by Day 4 and 5 of observation (Supplementary Figure 3A *middle* and *right panels*).

**Figure 4.**
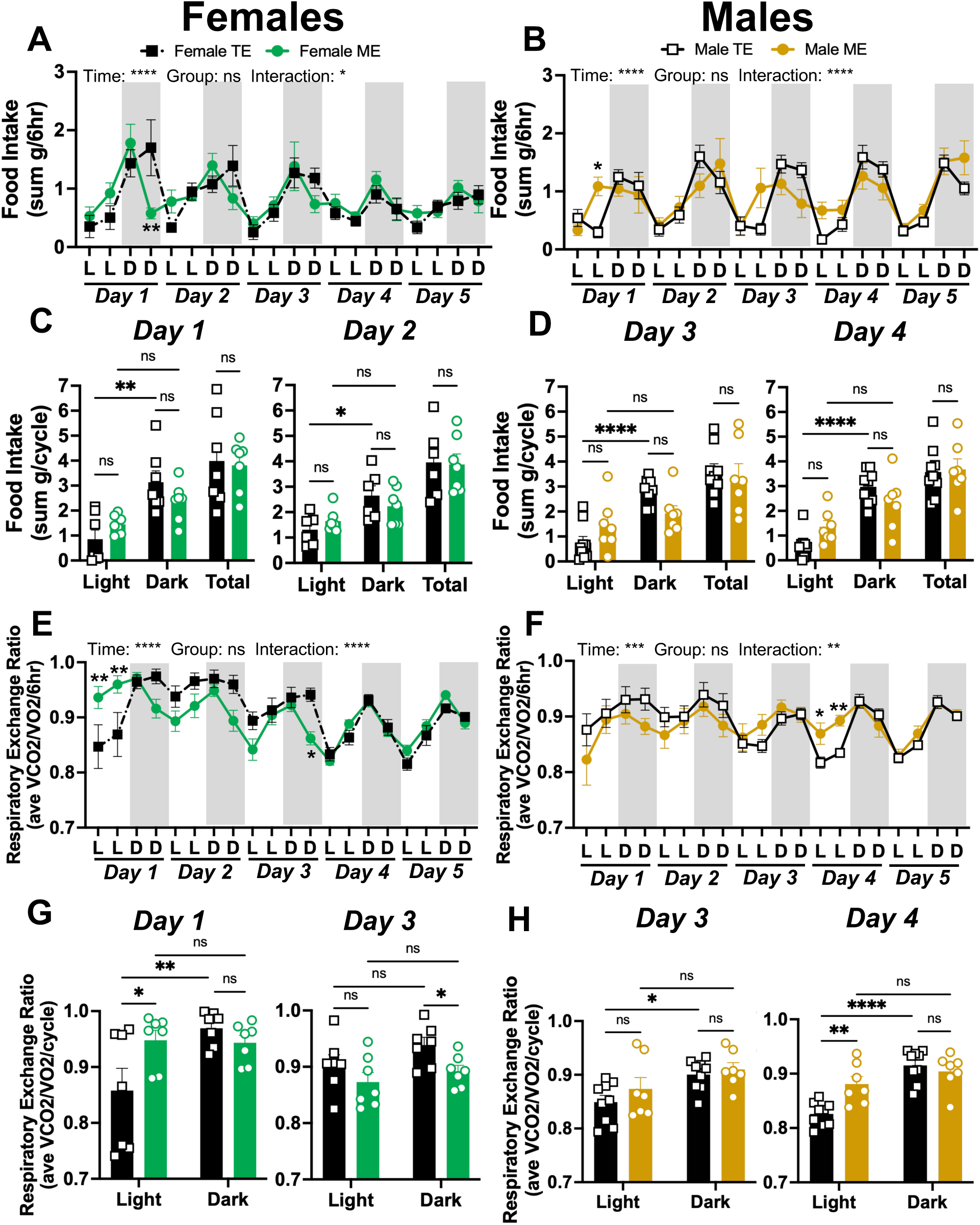
A single bout of mistimed eating disrupts the patterns of food intake and macronutrient utilization. (A) Sum of food intake every 6 hours across the light (L) and dark (D) cycles for 5 days after timed eating (TE) or mistimed eating (ME) in female animals. TE *n* = 7. ME *n =* 7. (B) Sum of food intake every 6 hours across the L and D cycles for 5 days after TE or ME in male animals. TE *n* = 9. ME *n =* 7. (C) The sum of food consumed during the light cycle, dark cycle, and total over 24 hours in female mice 1 (*left*) and 2 (*right*) days after TE or ME. TE *n* = 7. ME *n =* 7. (D) The sum of food consumed during the light cycle, dark cycle, and total over 24 hours in female mice 3 (*left*) and 4 (*right*) days after TE or ME. TE *n =*9. ME *n* = 7. (E) Average of respiratory exchange ratio (RER) every 6 hours across the L and D cycles for 5 days after TE or ME in female animals. TE *n* = 7. ME *n =* 7. (F) Average of RER every 6 hours across the L and D cycles for 5 days after TE or ME in male animals. TE *n* = 9. ME *n =* 7. (G) The average of RER during the light and dark cycle over 24 hours in female mice 1 (*left*) and 3 (*right*) days after TE or ME. TE *n* = 7. ME *n =* 7. (H) The average of RER during the light and dark cycle over 24 hours in female mice 3 (*left*) and 4 (*right*) days after TE or ME. TE *n =*9. ME *n* = 7. Values are means ± SEM. In Figures 4A, 5B, 5E, and 5F significance was determined by repeated measures 2-way ANOVA with Holm-Šídák’s post hoc test. In Figures 4C, 4D, 4G, and 4F, significance was determined by 2-way ANOVA with Tukey’s post hoc test. ns= no significance, *p<0.05, **p<0.01, ****p<0.0001.

ME disrupted food intake patterns in male animals for 4 days after ME (males Time x Group: *p=* 0.0086; Figure 4B). It took animals 5 days to recover from ME, as ME eliminated light:dark variation in food intake through Day 4 (Figure 4D *left* and *right panel;* Supplementary Figure 3B *left* and *middle panel*). ME animals did not fully regain light:dark variation in food intake until Day 5 (Supplementary Figure 3B *right panel*), at which point ME males consumed more total food (*p=* 0.0474; Supplementary Figure 3B *right panel*). Male TE animals retained daily patterns in food intake for the entire 5-day observation period (Figure 4B and D; Supplementary Figure 3B).

ME disrupted the daily patterns of macronutrient utilization in female animals for three days (females Time x Group: *p*< 0.0001, Figure 4E). Female ME animals were metabolically inflexible for 3 days after ME (Figure 4G *left panel;* Supplementary Figure 3C *left panel*). Furthermore, female ME animals displayed a preference for carbohydrate utilization during the light phase on Day 1 (females Day 1: *p=* 0.0120, Figure 4G), but decreased preference for carbohydrate utilization on Day 3 (females Day 3: *p=* 0.0396, Figure 4G *right panel*). However, both TE and ME abolished daily variation in RER on Day 2 and Day 3 indicating a general effect of the TE and ME paradigms on macronutrient utilization (Figure 4G *right panel*; Supplementary Figure 3C *left panel*).

Males exhibited prolonged disruption in macronutrient utilization in response to ME as the daily patterns of RER were altered by ME for 4 days (males Time x Group: *p=* 0.0390; Figure 4F). TE and ME both abolished daily variation in RER for 2 days (Supplementary Figure 3D *left* and *middle* panel).

However, ME animals remained metabolically inflexible for Day 3 (Figure 4H *left* panel) and Day 4 (Figure 4H *right* panel) post-ME and exhibited increased carbohydrate utilization during the light phase on Day 4 (males Day 4: *p=* 0.0012, Figure 4H *left* panel). TE animals recovered light:dark variation in RER on Day 3 (Figure 4H *left* panel), but ME animals did not recover until Day 5 (Supplementary Figure 3D *right* panel).

### A Single Bout of Mistimed Eating Disrupts the Daily Patterns of Rest

ME increased the amount of time spent resting over the 5-day observation period in females (Time x Group: *p=* 0.0015, Figure 5A). Like EE and wheel-running, disruptions in rest in ME females did not manifest until Day 3, Day 4, and Day 5 post-ME (Figure 5C; Supplementary Figure 4A). Specifically, ME increased time spent resting in the dark phase on Day 3 (*p=* 0.0020, Figure 5C *left panel*) and Day 4 (*p*< 0.0001, Figure 5C *right panel*), but increased time spent resting in the light phase on Day 5 (*p=* 0.0361, Figure 5C *bottom panel*). Despite these disruptions, light:dark variation in time spent resting was not impacted by ME (Figure 5A and C; Supplementary Figure 4A).

**Figure 5.**
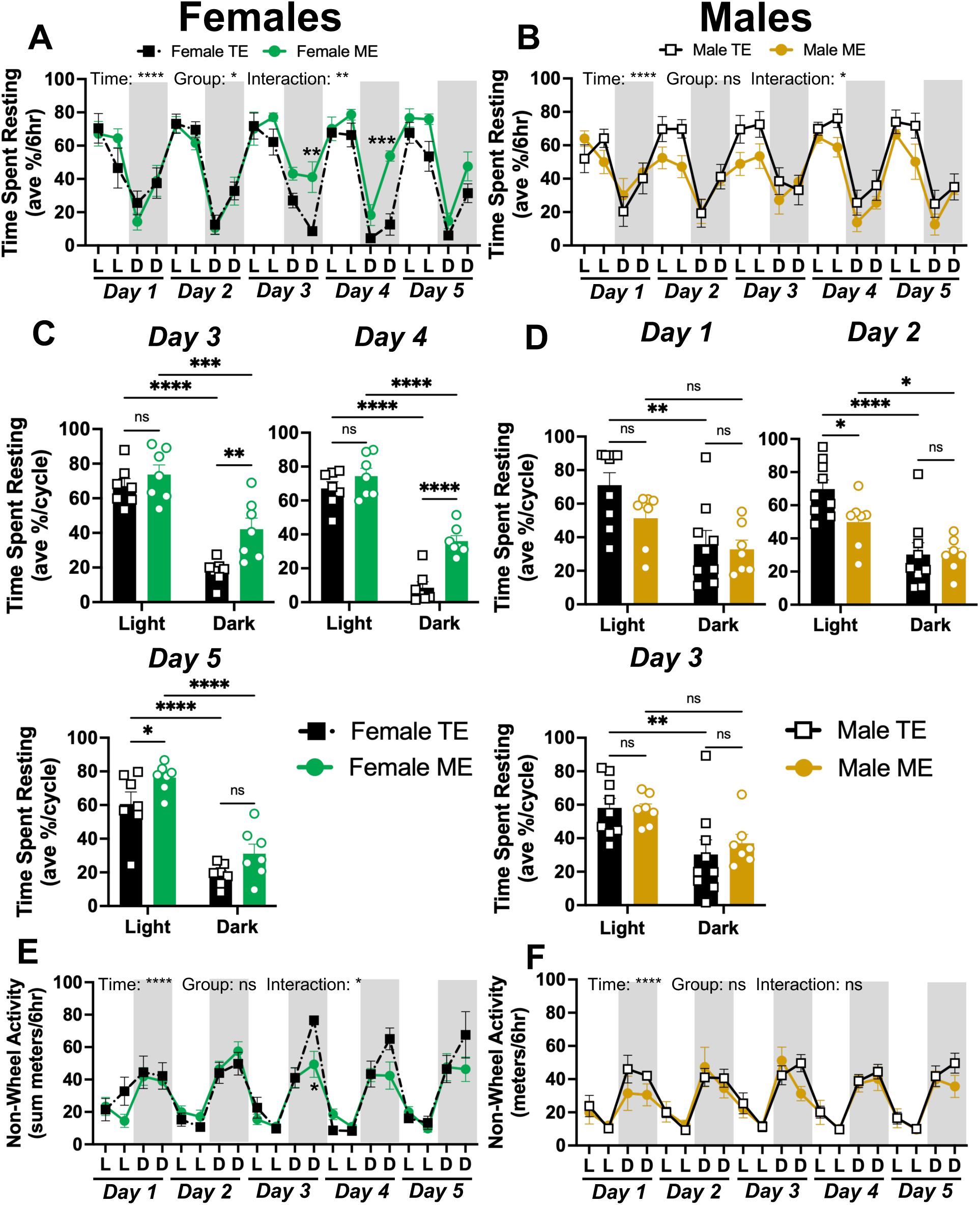
A single bout of mistimed eating disrupts daily resting patterns. (A) Average of percent of time spent resting every 6 hours across the light (L) and dark (D) cycles for 5 days after timed eating (TE) or mistimed eating (ME) in female animals. TE *n* = 7. ME *n =* 7. (B) Average of percent of time spent resting every 6 hours across the L and D cycles for 5 days after TE or ME in male animals. TE *n* = 9. ME *n =* 7. (C) The average percent of time spent resting in the light and dark cycle in female mice 3 (*left*), 4 (*right*), and 4 (*bottom*) days after TE or ME. TE *n =*7. ME *n* = 7. (D) The average percent of time spent resting in the light and dark cycle in male mice 1 (*left*), 4 2*right*), and 3 (*bottom*) days after TE or ME. TE *n =*9. ME *n* = 7. (E) Sum of non-wheel activity every 6 hours across the L and D cycles for 5 days after TE or ME in female animals. TE *n* = 7. ME *n =* 7. (B) Sum of non-wheel activity every 6 hours across the light (L) and dark (D) cycles for 5 days after TE or ME in male animals. TE *n* = 9. ME *n =* 7. Values are means ± SEM. In Figures 5A, 5B, 5E, and 5F significance was determined by repeated measures 2-way ANOVA with Holm-Šídák’s post hoc test. In Figures 5C and 5D significance was determined by 2-way ANOVA with Tukey’s post hoc test. ns= no significance, *p<0.05, **p<0.01, ***p<0.001, ****p<0.0001.

ME reduced the amount of time spent resting over the 5-day observation period in males (Time x Group: *p=* 0.0241, Figure 5B). ME abolished light:dark variation in resting in males on Day 1 (Figure 5D *left panel*). On Day 2, ME males exhibited reduced time spent resting in the light phase (*p=* 0.0256, Figure 5D *right panel*), but regained daily variation in resting (*p=* 0.0356, Figure 5D *right panel*). Again, on Day 3 ME caused light:dark variation in resting to be lost (Figure 5D *bottom panel*). However, by Days 4 and 5 ME males fully recovered light:dark variation in resting (Supplementary Figure 4B *left* and *right panel*).

Female and male TE animals retained daily patterns in time spent resting for the entire 5-day observation period (Figure 5A-D; Supplementary Figure 4A-B).

We hypothesized that ME-induced reduction in resting would be accompanied by an increase in general, non-wheel cage activity specifically during the light phase. Female ME animals exhibited a reduction in non-wheel cage activity. Specifically, ME induced a reduction in non-wheel activity in the late dark phase on Day 3 (p= 0.0323, Figure 5E), which corresponded with the increased time spent resting during the late dark phase on Day 3. We did not observe an effect of ME on non-wheel activity in male animals (male Time x Group: *p=* 0.3316, Figure 5F). Female and male TE animals retained daily patterns in non-wheel activity for the entire 5-day observation period (Figure 5E-F).

### A Single Bout of Mistimed Eating Does Not Impact Gross Anatomy or Physiology

On Day 6 at ZT3, TE and ME animals were euthanized to assess the gross anatomy and physiology of recently fed animals while avoiding the potential confounding variable of lights on at ZT0. ME had no effect on final body weight, non-fasting blood glucose, normalized liver weight, normalized gWAT weight, or normalized iWAT weight in females or males (Table 1–2).

## DISCUSSION

This study found that a single bout of mistimed eating is sufficient to cause significant disruptions in metabolic and behavioral rhythms for several days in female and male mice. The ME paradigm disrupted EE and wheel running activity while altering light:dark variation in food intake and disrupting macronutrient utilization. Furthermore, ME changed the amount of time spent resting for several days after the eating event. The TE paradigm disrupted macronutrient utilization for two days but otherwise had no effect on metabolism or behavior. Our results demonstrate that acute ME has lasting impacts on daily rhythmicity and is an understudied but critical source of circadian disruption. Ultimately, this work suggests that even intermittent chrono-disruptions are physiologically consequential, and that a single mistimed meal can promote metabolic dysfunction.

EE serves as an important metabolic readout due to its systemic influence on the body (19). It is particularly relevant to the cardiovascular system, as aortic blood flow carries the necessary nutrients to all other tissues in the body, allowing for energy use (20). The cardiovascular system also contains oscillatory changes in factors such as blood pressure, heart rate, and hormonal regulation exhibiting time-of-day variations. Perturbations in these rhythms have been linked to increased occurrence and exacerbation of obesity and hypertension in mice and humans (21). Research strongly suggests that exercise can have immediate effects on EE, with increased total EE being linked to protective mechanisms for cardiometabolic outcomes (22). A single bout of ME significantly reduced the total EE of male and female mice. Circadian variability between L:D cycles was not altered in the females but was abolished in the males. Thus, ME mice spent less energy during their active phase compared to TE mice. This disruption is likely explained by changes in high-intensity, wheel-running activity, as following the ME bout, mice spent less time engaging in high-intensity activity and expended less energy during their active phase. Consequently, the correlation in reduced EE and reduced activity seen after one bout of ME may worsen cardiometabolic outcomes, though further investigation is necessary. Currently, some therapies of obesity and type 2 diabetes aim to leverage EE patterns in the development of new treatments (23).

A single bout of ME abolished circadian variation in RER and altered food intake patterns in male and female mice. Though total food consumption was not significantly different, ME mice consumed more food during rest periods and less food during active periods compared to TE mice. This disruption likely explains the transient metabolic inflexibility observed in ME animals. Indeed, ME mice exhibited reduced L:D variation in RER, indicating an inability to efficiently switch between carbohydrate and lipid utilization during feeding and fasting states, respectively.

Metabolic flexibility is a crucial health parameter as it indicates an organism’s ability to access and use different energy stores. It provides information on how well the body can tolerate diverse metabolic demands and is linked to pre- and post-prandial glucose and insulin responses (24). In healthy individuals, a carbohydrate-heavy meal will drive the RER toward a value of 1, indicating carbohydrate usage over lipids, while a high-fat meal or prolonged fast will drive the RER toward 0.7, indicating lipid utilization over carbohydrates (25). The inability to effectively respond to energy demands and utilize macronutrients can negatively affect glucose handling, insulin sensitivity, dyslipidemia, and ectopic fat storage, which ultimately leads to chronic metabolic diseases such as obesity, type 2 diabetes, and nonalcoholic fatty liver disease (26–30). Therefore, ME-induced disruptions in RER and food intake may precede advanced metabolic disease. However, determining this requires further testing.

A single bout of ME significantly impacted the amount of time spent resting. Interestingly, resting patterns exhibited a sex-specific effect. Female mice spent more overall time resting and did not show abolished circadian variability. Male mice spent less overall time resting and exhibited intermittent abolished circadian variability. Thus, female ME mice spent more time resting during their active phase and male ME mice spent less time resting during their rest phase compared to TE mice. This disruption was likely linked to changes in low-intensity, non-wheel running activity. Following the ME bout, female mice were less engaged in low-intensity activity and instead showed a preference for resting. However, male mice exhibited similarly decreased engagement in low-intensity activity despite their decreased rest during the resting phase, suggesting that there may be a sex-specific effect of ME on general, non-wheel activity.

Resting is an essential circadian behavior in mammals, broadly including periods of inactivity or sleep. The balance between activity and rest allows mammals to regulate energy expenditure, which becomes relevant in behavioral considerations—survival, foraging, etc. Sleep is crucial for the removal of waste and toxins from the central nervous system, and under normal circadian rhythms, influences a multitude of hormonal signaling pathways, playing an important role in weight regulation, appetite, and other metabolic processes (31,32). Research has shown that disturbances in sleep patterns induce weight gain and increase the risk of diabetes, i.e., working through upregulated appetite, decreased EE, and altered glucose pathways (33). However, sleep is a unique outcome in that it is often used to diagnose depression, anxiety, and other mood disorders (34). Thus, resting patterns influence not just metabolic health, but mental health as well, with studies showing that late-night eating affects serotonin and dopamine rhythms, causing emotional disturbances and increasing the risk of mood disorders (35). In addition, the effects of disrupted sleep rhythms can converge with energy expenditure and substrate utilization disruptions, with studies revealing relationships to hypertension and heart disease (36). While we are conducting a separate study to determine the effects of acute ME on depressive-type behavior, further characterization of resting and sleep as ME outcomes requires investigation.

Our data also revealed sex differences in the timelines that the metabolic and behavioral effects of ME manifest on. Specifically, females are more susceptible to delayed disrupted effects while males are more susceptible to early disruptive effects of ME. Sex differences in metabolic and behavioral parameters have been well characterized in the literature. Female sex hormones are typically protective against metabolic disease, while circadian disruption has a more pronounced effect on sleep, cognition, and mood in females compared to males (37,38). These differences are of considerable interest given the influence of sex hormones within the neural structures driving circadian system timing (39). Studies have primarily established that the amplitude of disruption in sleep and cognitive effects were more pronounced in women compared to men, with differences persisting down to the molecular level; average levels of leptin, a satiety hormone, decreased in females and increased in males, along with increased ghrelin expression, a hunger hormone, in females, with no considerable difference in males (40). Despite these hormonal changes, males actually reported increased food cravings, implicating activity of hedonic appetite pathways (40); an interesting finding, as the broader literature suggests that females are more prone to binge eating (41,42). Additionally, many physiological processes implicated in metabolic disease have been linked to changes in the gut microbiome (43). Conditions altering the composition of the microbiota (such as a high-fat diet) can affect circadian clock gene expression, attenuating genes associated with liver lipogenesis, mitochondrial activity, and insulin sensitivity in females (43). However, a direct comparison of female and male responses to a single bout of ME requires further investigation.

Although we did not investigate the molecular mechanisms involved in the metabolic and behavioral effects of acute ME, we believe that internal desynchrony between the brain and the liver may play an important role. The liver is particularly vulnerable to ME-induced disruptions while the SCN remains entrained to light cycles, and internal desynchrony between the liver and SCN clocks exacerbates metabolic disease outcomes (30,31,44–48). Thus, we can speculate that desynchrony between the SCN and the liver underlies the metabolic effects of acute ME. It is likely that the liver requires several days to re-entrain ad libitum feeding after a single bout of ME, and this delayed realignment between the central and peripheral clock disrupts metabolic and behavioral rhythms. However, this hypothesis requires additional testing. Additionally, the impact of diet composition on the effects of acute ME on metabolism and behavior are not well understood. This study utilized a normal rodent chow diet, but a high-fat diet (HFD) is shown to affect multiple facets of circadian rhythmicity, from insulin and glucose rhythms to liver-specific rhythms (49). The effect of a HFD, or varying diet constitutions, may exaggerate the effect of ME or exacerbate existing metabolic dysfunction, but is beyond the scope of the current work. Finally, it is essential to design future research efforts investigating whether repeated bouts of acute ME without sufficient recovery from the preceding bout enable accumulation of chrono-metabolic disruption, ultimately resulting in exacerbated diseased states.

A major limitation of our study is that the experimental design requires mice to undergo a 24hr extended fast. This is necessary for our research question as mice will not consume food in their resting phase unless fasted. We could not utilize a 12hr fast because the ME and TE mice would be refed under different metabolic conditions; since mice are nocturnal, they naturally fast during the light (rest) phase. In this scenario, ME mice would have begun a 12hr fast at ZT13 and missed their natural fasting window, while TE mice would have begun a 12hr fast at ZT1, during their natural fasting window. Therefore, ME mice would have effectively been fasted for 24hrs while TE mice would have only fasted for 12hrs. Thus, to adequately compare a single bout of ME to a single bout of TE, a 24hr extended fast was necessary.

However, we recognize that prolonged fasting can induce alterations in EE and RER in mammals (50,51). Indeed, we observed in our data that TE abolished RER rhythms in males for two days. Conversely, TE did not impact the other metabolic or behavioral outcomes, indicating that the main effect of our fasting/feeding paradigm was due to mistimed eating rather than the fast.

Overall, our data demonstrates that a single bout of ME is sufficient to induce persistent disruptions in metabolic and behavioral rhythms that last several days beyond the feeding event. These findings are supported by the plethora of research showing that chronic ME significantly alters metabolic rhythms, but challenges the prevailing assumption that acute chrono-disruption is physiologically inconsequential (52–55). Our work suggests that acute ME is an unrecognized source of circadian disruption capable of inducing lasting metabolic consequences, providing a critical avenue of future investigation.

## DATA AVAILABILITY

All data included in this publication will be made available upon request to the corresponding author.

## SUPPLEMENTAL MATERIAL

Supplemental Figs. S1-S4

## ACKNOWLEDGMENTS

The authors would like to thank the mice used in this study for giving their lives to advance our understanding in how mistimed eating impacts metabolic and behavioral health. We would like to thank the GWU Office of Animal Research for the exemplary care they provided our animals.

## AUTHOR CONTRIBUTIONS

Identify which authors participated in the research: Conceived and designed research-SD, MY, END, LNW, performed experiments-SD, MY, END, LNW, analyzed data-LNW, interpreted results of experiments-SD, LNW, prepared figures-LNW, drafted manuscript-SD, LNW, edited and revised manuscript-SD, MY, END, LNW, approved final version of manuscript-SD, MY, END, LNW.

---

**Supplementary Figure 1.**
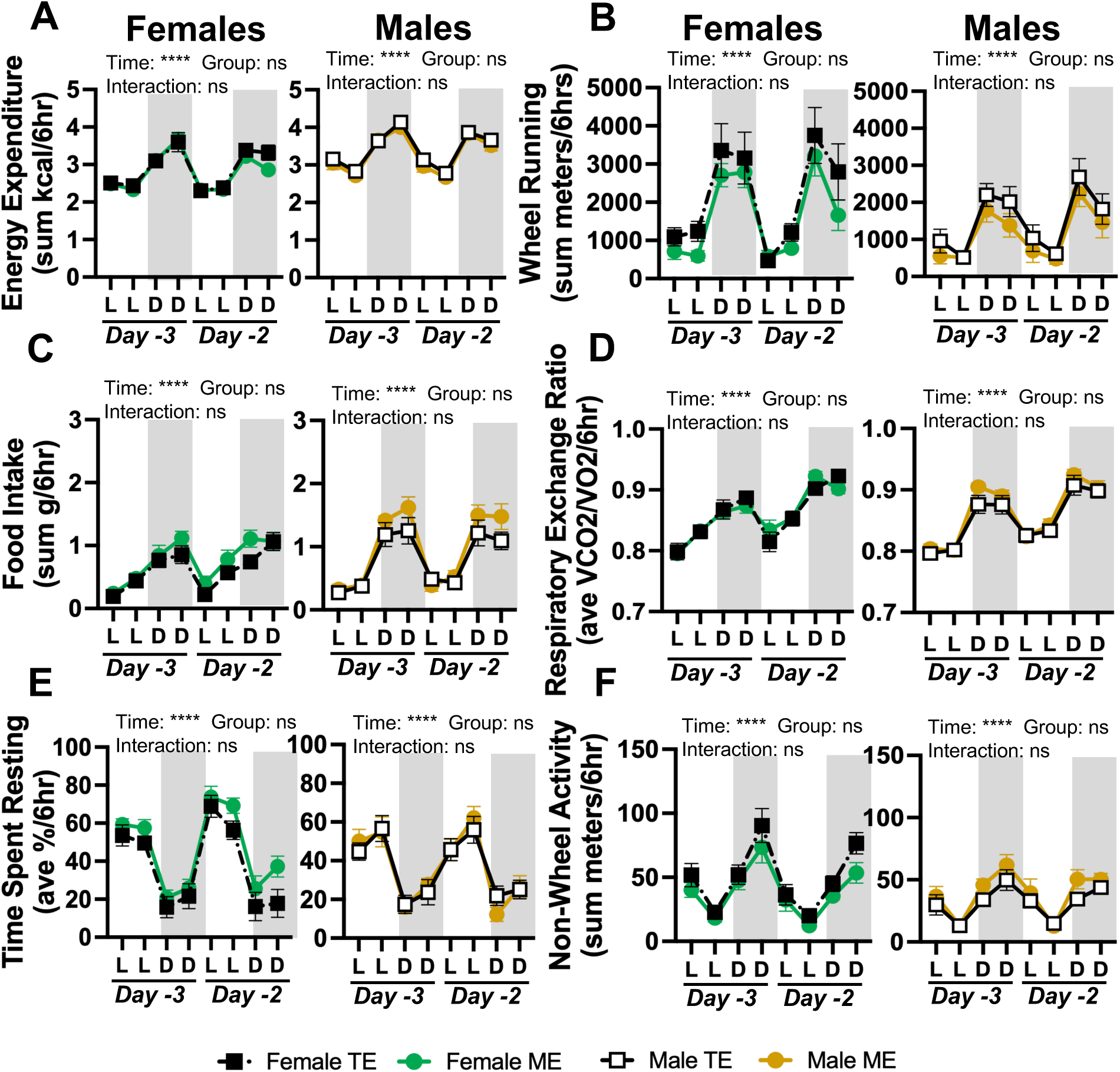
Baseline metabolic and behavioral measurements in male and female mice. (A) Sum of energy expenditure every 6 hours during the two days of baseline measurements in female (*left*) and male (*right*) mice. Female timed eating (TE) *n* = 8. Female mistimed eating (ME) *n* = 8. Male TE *n* = 11. Male ME *n* = 11. (B) Sum of wheel-running every 6 hours during the two days of baseline measurements in female (*left*) and male (*right*) mice. Female TE *n* = 7. Female ME *n* = 7. Male TE *n* = 9. Male ME *n* = 10. (C) Sum of food intake every 6 hours during the two days of baseline measurements in female (*left*) and male (*right*) mice. Female TE *n* = 8. Female ME *n* = 8. Male TE *n* = 11. Male ME *n* = 11. (D) Average of respiratory exchange ratio every 6 hours during the two days of baseline measurements in female (*left*) and male (*right*) mice. Female TE *n* = 8. Female ME *n* = 8. Male TE *n* = 11. Male ME *n* = 11. (E) Average of percent of time spent resting every 6 hours during the two days of baseline measurements in female (*left*) and male (*right*) mice. Female TE *n* = 7. Female ME *n* = 8. Male TE *n* = 11. Male ME *n* = 11. (F) Sum of non-wheel activity every 6 hours during the two days of baseline measurements in female (*left*) and male (*right*) mice. Female TE *n* = 8. Female ME *n* = 8. Male TE *n* = 11. Male ME *n* = 11. Values are means ± SEM. Significance was determined by repeated measures 2-way ANOVA with Holm-Šídák’s post hoc test. ns= no significance, ****p<0.0001.

**Supplementary Figure 2.**
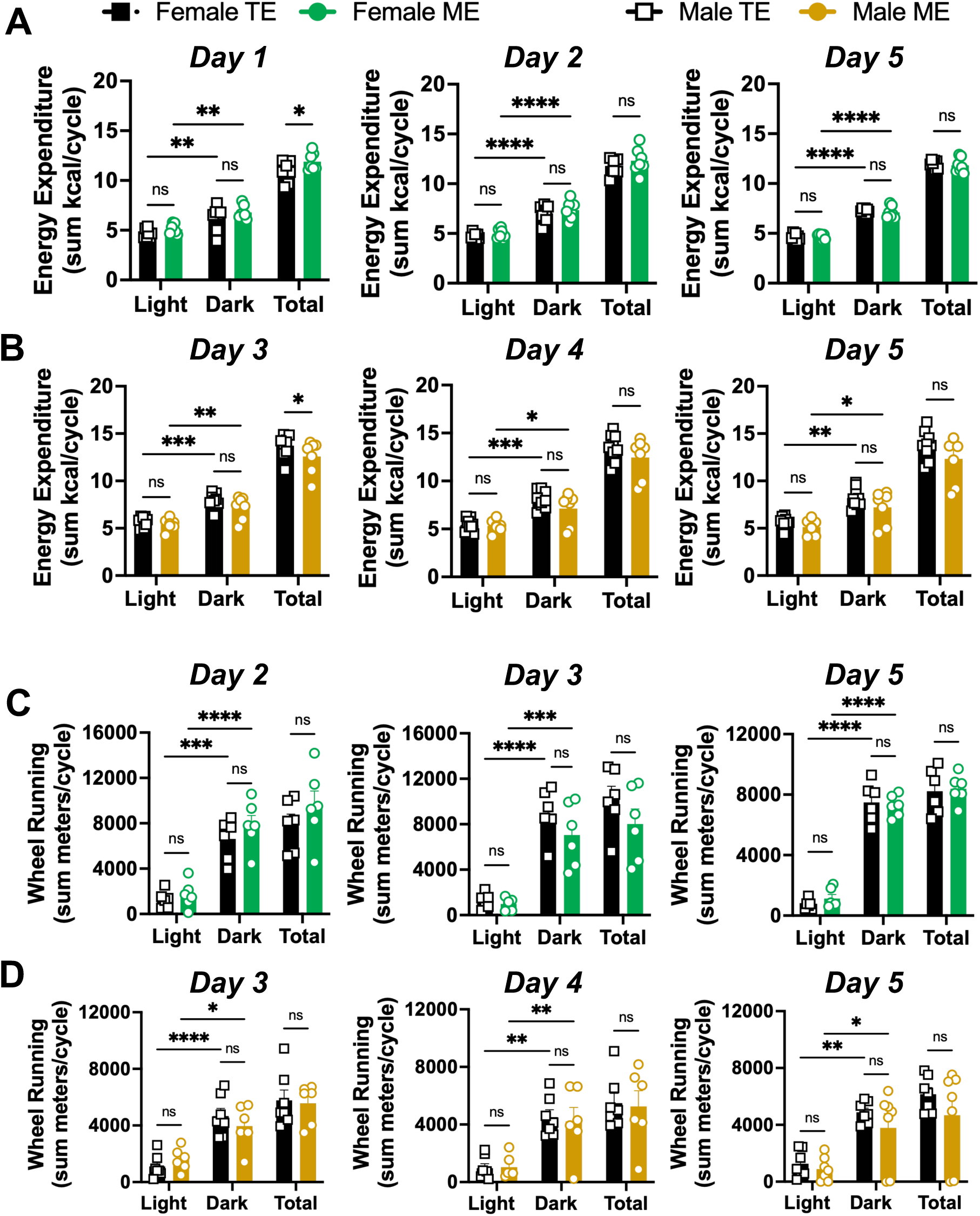
Additional days of energy expenditure and wheel-running cycle analysis. A) The sum of energy expended during the light cycle, dark cycle, and total over 24 hours in female mice 1 (*left*), 2 (*middle*), and 5 (*right*) days after timed eating (TE) or mistimed eating (ME). TE *n* = 7. ME *n =* 7. (B) The sum of energy expended during the light cycle, dark cycle, and total over 24 hours in male mice 3 (*left*), 4 (*middle*), and 5 (*right*) days after TE or ME. TE *n* = 9. ME *n =* 7. (C) The sum of meters run on the wheel during the light cycle, dark cycle, and total over 24 hours in female mice 2 (*left*), 3 (*middle*), and 5 (*right*) days after TE or ME. TE *n* = 7. ME *n =* 7. (D) The sum of energy expended during the light cycle, dark cycle, and total over 24 hours in male mice 3 (*left*), 4 (*middle*), and 5 (*right*) days after TE or ME. TE *n* = 9. ME *n =* 7. Values are means ± SEM. Significance was determined by 2-way ANOVA with Tukey’s post hoc test. ns= no significance, *p<0.05, **p<0.01, ***p<0.001, ****p<0.0001.

**Supplementary Figure 3.**
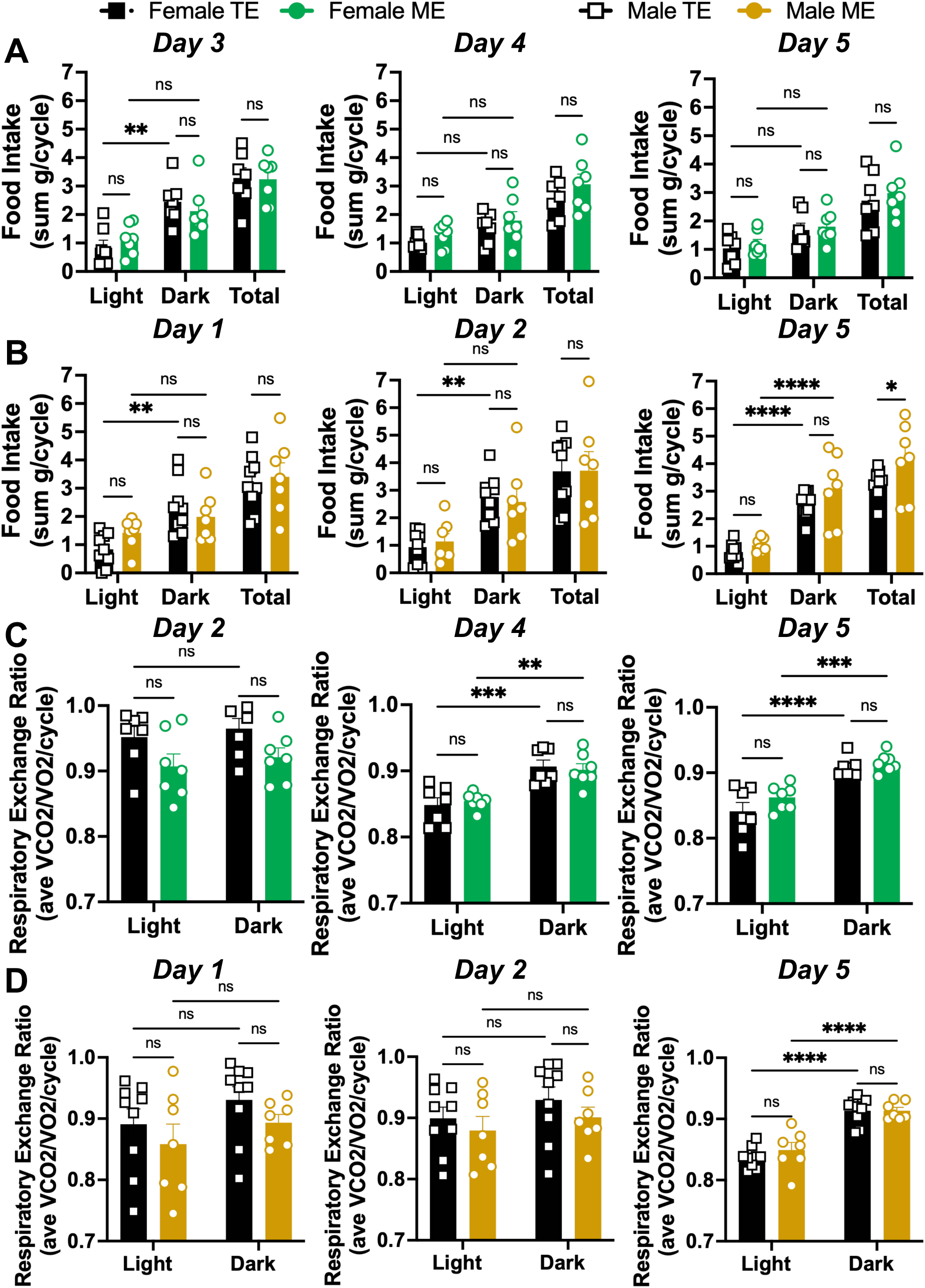
Additional days of food intake and respiratory exchange ratio cycle analysis. (A) The sum of food consumed during the light cycle, dark cycle, and total over 24 hours in female mice 3 (*left*), 4 (*middle*), and 5 (*right*) days after timed eating (TE) or mistimed eating (ME). TE *n* = 7. ME *n =* 7. (B) The sum of food consumed during the light cycle, dark cycle, and total over 24 hours in male mice 1 (*left*), 2 (*middle*), and 5 (*right*) days after TE or ME. TE *n* = 9. ME *n =* 7. (C) The average respiratory exchange ratio (RER) in the light and dark cycle in female mice 2 (*left*), 4 (*middle*), and 5 (*right*) days after TE or ME. TE *n* = 7. ME *n =* 7. (D) The average respiratory exchange ratio (RER) in the light and dark cycle in male mice 21(*left*), 2 (*middle*), and 5 (*right*) days after TE or ME. TE *n* = 9. ME *n =* 7. Values are means ± SEM. Significance was determined by 2-way ANOVA with Tukey’s post hoc test. ns= no significance, *p<0.05, **p<0.01, ***p<0.001, ****p<0.0001.

**Supplementary Figure 4.**
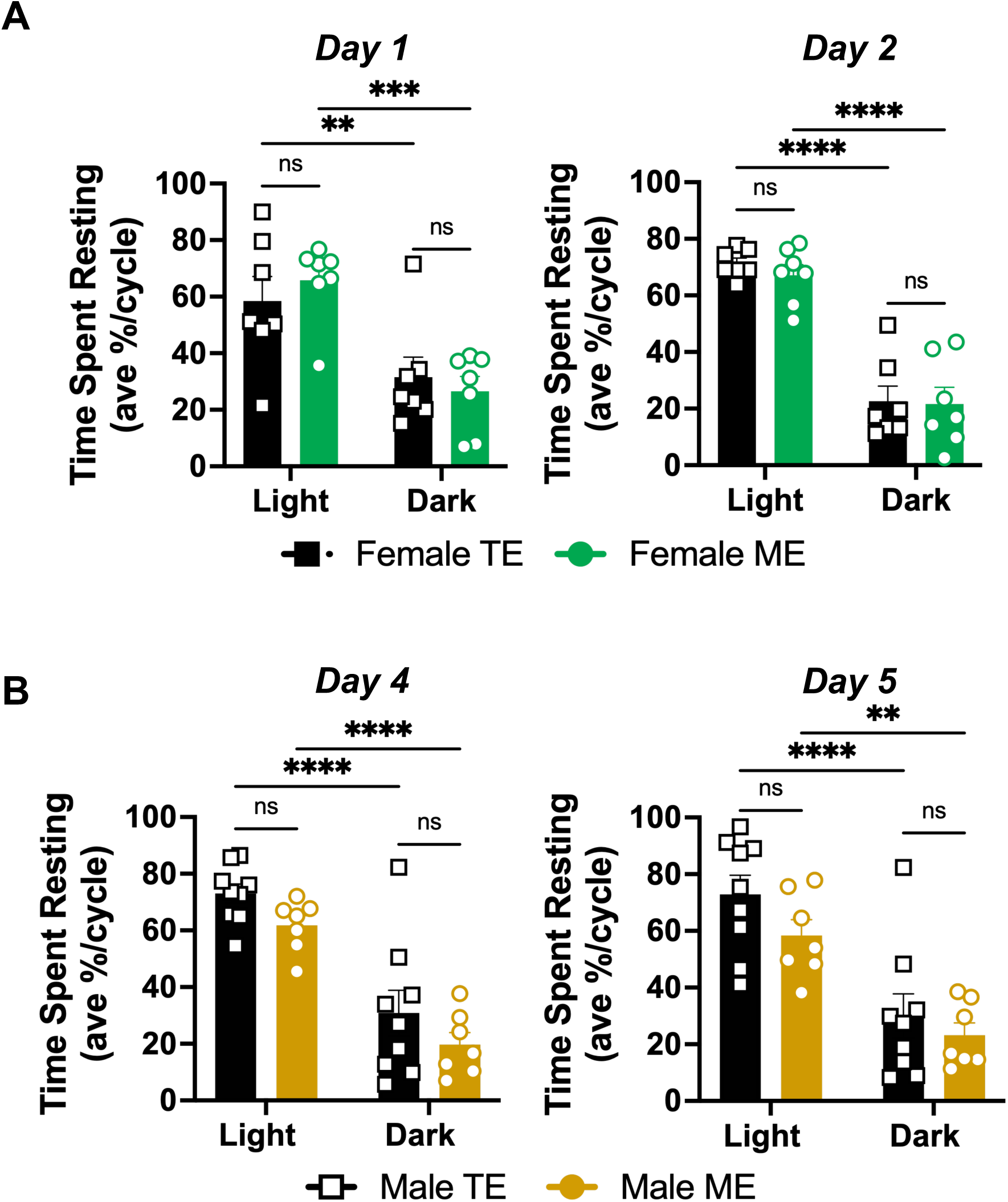
Additional days of time spent resting cycle analysis. (A) The average percent of time spent resting in the light and dark cycle in female mice 1 (*left*) and 2 (*right*) days after timed eating (TE) or mistimed eating (ME). TE *n =*7. ME *n* = 7. (B) Light and dark cycle analysis of time spent resting in male mice 4 (*left*) and 5 (*right*) days after TE or ME. TE *n =*9. ME *n* = 7. Values are means ± SEM. Significance was determined by 2-way ANOVA with Tukey’s post hoc test. ns= no significance, **p<0.01, ***p<0.001, ****p<0.0001.

## GLOSSARY

ME: Mistimed Eating
TE: Timed Eating
ZT: Zeitgeber Time
EE: Energy Expenditure
RER: Respiratory Exchange Ratio

